# Reconstitution of ultrawide DNA origami pores in liposomes for transmembrane transport of macromolecules

**DOI:** 10.1101/2021.02.24.432733

**Authors:** Alessio Fragasso, Nicola De Franceschi, Pierre Stömmer, Eli O. van der Sluis, Hendrik Dietz, Cees Dekker

**Affiliations:** Department of Bionanoscience, Kavli Institute of Nanoscience, Delft University of Technology, Van der Maasweg 9, 2629 HZ Delft, The Netherlands; Physik Department, Technische Universität München, Am Coulombwall 4a, Garching bei München D-85748, Germany

**Keywords:** DNA origami, nanopores, liposomes, transmembrane transport, cDICE

## Abstract

Molecular traffic across lipid membranes is a vital process in cell biology that involves specialized biological pores with a great variety of pore diameters, from fractions of a nanometer to >30 nm. Creating artificial membrane pores covering similar size and complexity will aid the understanding of transmembrane molecular transport in cells, while artificial pores are also a necessary ingredient for synthetic cells. Here, we report the construction of DNA origami nanopores that have an inner diameter as large as 30 nm. We developed new methods to successfully insert these ultrawide pores into the lipid membrane of giant unilamellar vesicles (GUVs) by administering the pores concomitantly with vesicle formation in an inverted-emulsion cDICE technique. The reconstituted pores permit the transmembrane diffusion of large macromolecules such as folded proteins, which demonstrates the formation of large membrane-spanning open pores. The pores are size selective as dextran molecules with a diameter up to 22 nm can traverse the pores, whereas larger dextran molecules are blocked. By FRAP measurements and modelling of the GFP influx rate, we find that up to hundreds of pores can be functionally reconstituted into a single GUV. Our technique bears great potential for applications across different fields from biomimetics, synthetic biology, to drug delivery.

## Introduction

In the last decade, advancements in the field of DNA nanotechnology have enabled the fabrication of a great variety of ‘DNA origami’ nanostructures^1,2^, including transmembrane structures that resemble the biological pores found in cells^3^. Drawing inspiration from protein pores such as alpha-hemolysin^4^, artificial pores have been engineered with DNA origami^1,2^ that can insert into lipid bilayers and allow for transmembrane diffusion of ions^5,6^ and small molecules such as fluorophores^7,8^, DNA oligomers^5,7^, short PEG^9^, and dextran^8,10^. To favor partitioning into the lipid bilayer, the usual strategy relies on the chemical modification of the outer nanopore surface with hydrophobic groups, e.g. cholesterols, that anchor and stabilize the strongly hydrophilic DNA-based nanopores into the lipid membrane^3^. As DNA origami structures can be designed to virtually any shape or size^11^ up to the dimensions comparable to those of viruses^12^, the approach bears great potential for various applications, from recapitulating the function of complex enzymes like flippase^13^, to mimicking large protein transporters such as the nuclear pore complex^14,15^.

However, so far only small pores with an inner diameter of a few nm have been realized^8,10^, which is because wider pores are increasingly difficult to insert into a membrane. A commonly used strategy to reconstitute pores into a bilayer relies on spontaneous insertion into a preformed lipid membrane^3^. As this method relies on transient membrane instabilities^16^, it is becoming increasingly ineffective for larger size objects. This is because the work required to open up a hole in a membrane grows quadratically with the diameter of the target object^6,8,17^, and spontaneous insertion occurs in a Boltzmann weighted fashion, thus exponentially decreasing the chances of success for larger objects. Scaling up the size of artificial pores thus requires an alternative insertion process that can circumvent the barriers associated with spontaneous insertion.

Here, we overcome these challenges by employing a continuous droplet interface crossing encapsulation (cDICE) technique^18^ to incorporate newly designed ultrawide (~30 nm inner diameter, ~55 nm outer diameter) DNA origami pores into the membrane of giant unilamellar vesicles (GUVs). As we show below, by administering the pores concomitantly with vesicle formation, the pores can be localized efficiently at the membrane. To demonstrate the transmembrane transport through the pores, we measured the influx of multiple fluorescent macromolecules with different sizes using confocal microscopy. We find that the origami pores indeed provide open portals for transport that support the molecule traffic of extremely large (22nm) molecular structures, consistent with the 30 nm inner diameter of the pores. Using fluorescence recovery after photobleaching (FRAP), we estimate that up to hundreds of pores can be functionally reconstituted in a single liposome. These ultrawide pores represent an exciting tool for various scopes, from the mimicking of large biological pores such as the nuclear pore complex, to applications in synthetic biology and drug delivery.

## Results

### Design and assembly of ultrawide DNA origami pores

Using DNA origami folding of a template strand^1,2^, we designed and assembled a rigid octagonal ring-like structure, see Fig.1. A multi-layer DNA origami object was designed in square-lattice helical packing and folded from a 7560 bases long scaffold single strand with 240 individual oligonucleotide single strands, arranged in a 4 × 4 double-helical pattern. It was designed to form a closed-loop octagonal shape, formed by 8 corner design motifs^19^ with single-stranded poly-T strands at the corner sites. Deletions were introduced every 32 bases to correct for global twist^20^. The origami pore was designed with a nominal outer diameter of 55 nm, an inner diameter of 35 nm, and a height of 10 nm (Fig.1a) which represents the channel length. The design includes 48 single stranded handles that were distributed evenly on the interior surface facing the central cavity, as sites for future functionalization purposes (Fig.S15). The octagon furthermore included 32 cholesterol modifications on the outer surface that were arranged evenly with a ~5.5 nm horizontal spacing and ~3 nm vertical distance between neighboring cholesterols (Fig.1a). An asymmetric feature was added (Fig.1b) consisting of four additional double helices on top of the octagon, arranged in a 2 × 2 double helical pattern on the top side of the octagon, spanning over 3 corners. The octagon further features 12 Atto647N dyes for fluorescence imaging, one at each corner on the bottom side of the octagon and four more on both ends of the asymmetric feature (Fig. 1a). We iteratively refined the octagon design using gel-electrophoretic mobility analysis as read-out, in order to minimize the occurrence of higher order aggregates and improve the folding quality of the octagon object (Figs.S3-S10).

**Figure 1.**
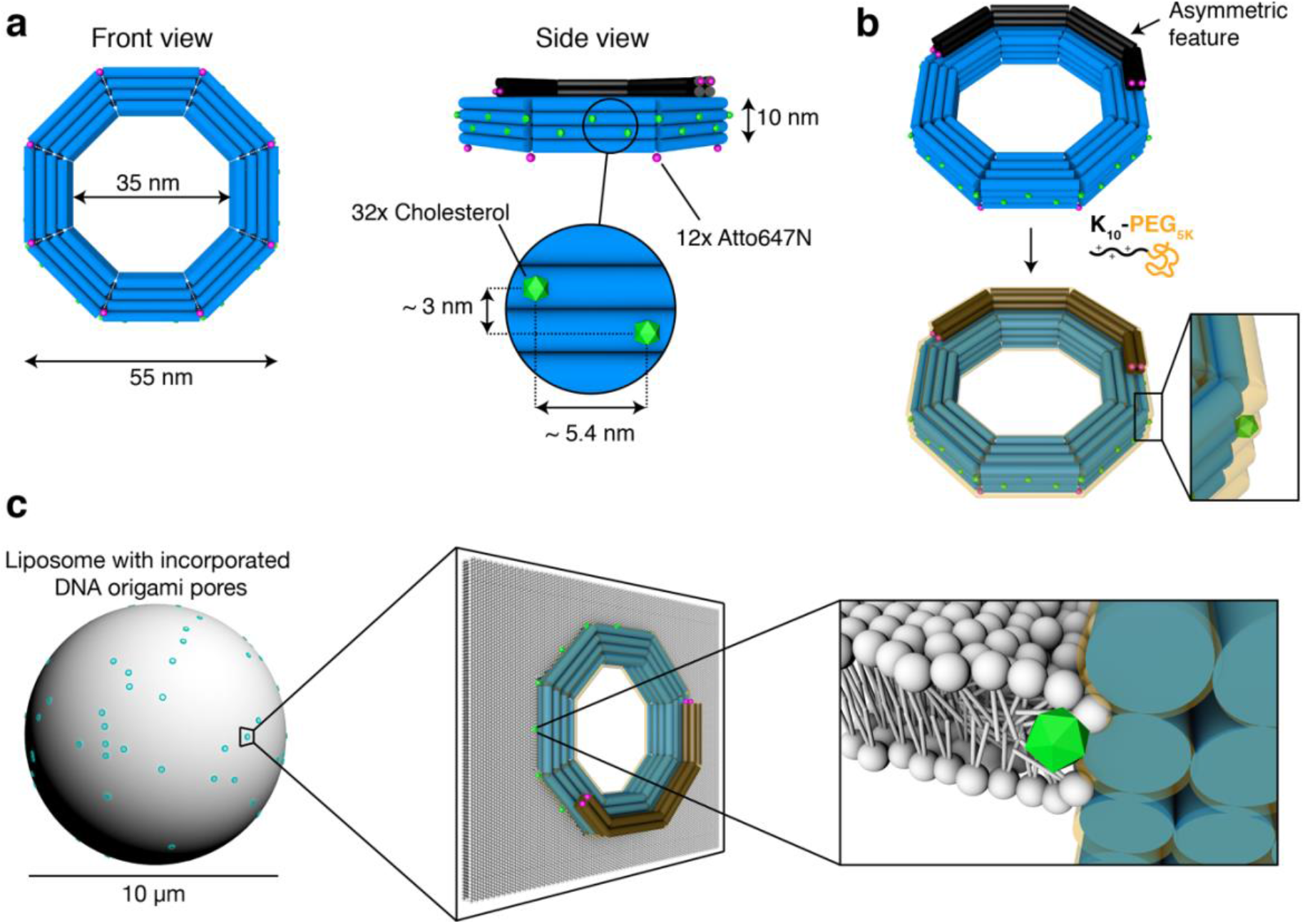
DNA origami pore design and incorporation approach. **a,** DNA origami pore in front (left) and side view (right). Blue cylinders represent DNA double helices. The outer diameter is 55 nm, the inner diameter is 35 nm. Green objects along the outer side of the DNA origami pore indicate the 32 cholesterol modifications. They vertically span over a width of ~3 nm (purposely slightly less than the hydrophobic thickness of ~3.7 nm of a DOPC-lipid bilayer^36^). Black cylinders represent an asymmetric feature that was introduced for class averaging of single particles from TEM. Violet spots along the outer surface of the DNA origami pore indicate the positions of 12 Atto647N dye modifications. The pore is designed with a height of 10 nm. **b,** Schematic representation of K_10_-PEG_5K_ coating of the origami pore. **c,** Schematic representation of the DNA origami pore incorporated into the lipid bilayer of a GUV. Left: a GUV with pores incorporated in the lipid bilayer; middle: zoom-in on one of the incorporated origami pores resulting in a 30 nm diameter hole; right: a further zoom-in showing individual lipid molecules of the bilayer; white spheres represent lipid heads, white strokes represent lipid tails, the green object represents a cholesterol.

The origami pore was coated with K_10_-PEG_5K_ molecules, which consist of 10 positively charged lysine amino acids (K) linked to a short polyethylene glycol chain (PEG, 5kDa) (Fig. 1B) as described in Ponnuswamy et al.^21^ This coating serves two purposes: (i) it stabilizes the octagon in low-ionic strength environments, where uncoated DNA origami structures would otherwise disassemble; and (ii) it prevents the cholesterol-modified origami pores from aggregating in bulk. Figure 1c shows a schematic illustration of the incorporation of origami pores into the lipid membrane of a liposome (left). The zoom-ins highlight the interaction of the cholesterol-modified origami pores with the lipid molecules.

Characterization of the proper folding and verification of the intended dimensions of the structure were performed using negative-stain TEM imaging and subsequent class averaging of single particles (Figure 2). Figure 2a shows a high-pass-filtered image of a typical field of view with several octagon pores. The overall shape of the majority of the particles match the intended design, without obvious defects, in which 8 straight edges and 8 corners form an octagonal ring-like structure with a ~35 nm inner diameter and a ~11 nm thickness. Most of the structures are oriented in a flat orientation on the surface, while a few can be seen in their side view.

**Figure 2.**
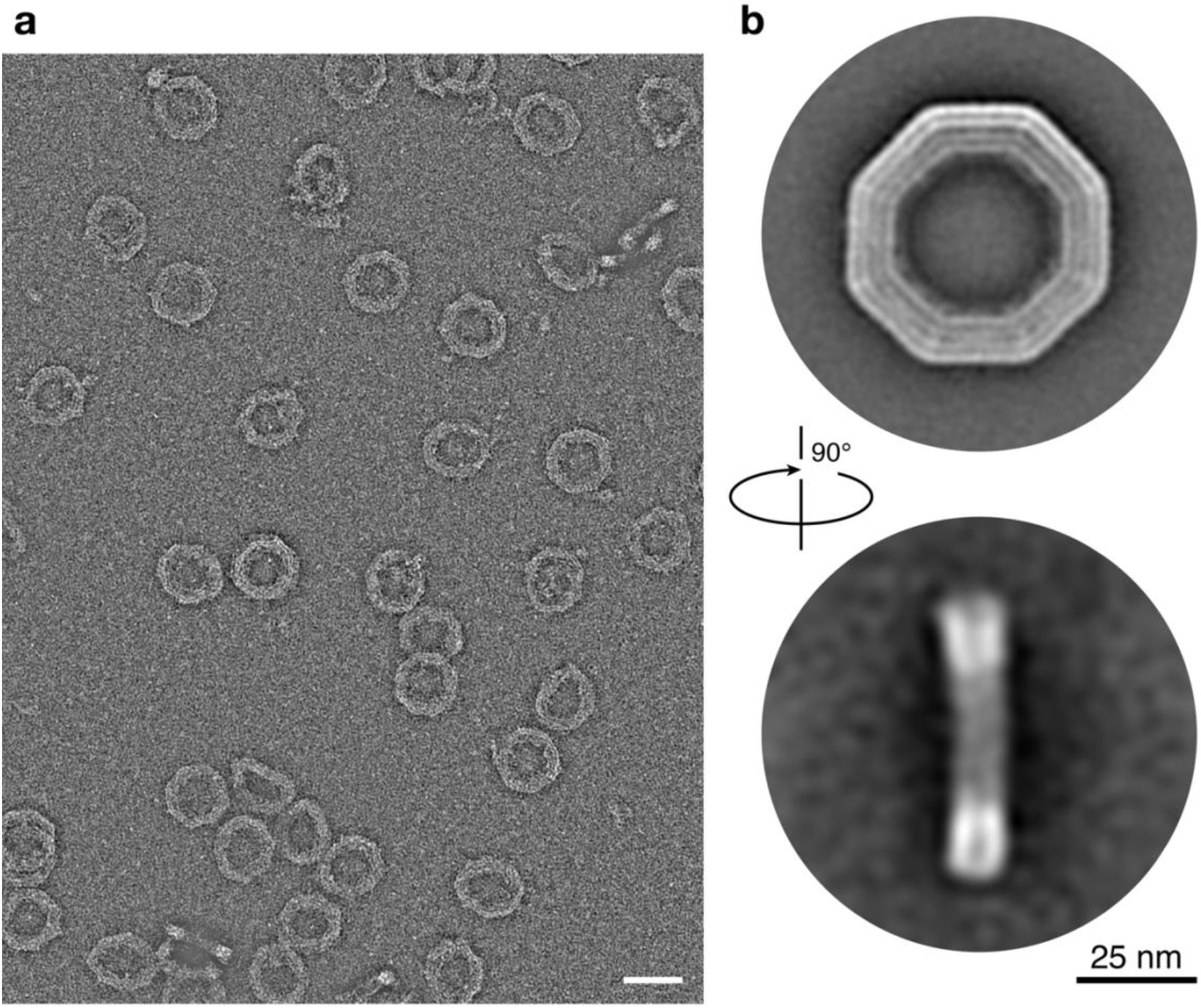
Negative-staining TEM and class averaging of the DNA origami pores. **a,** Typical field of view of a negative-stained DNA-origami sample. The pores in this image were prepared as described in the Methods section. The image is high-pass filtered with a radius of 20 nm and auto leveled in Photoshop. Scale bar: 50 nm. **b,** Top: Front view class average of N=682 single particles (Fig.S12, left). Bottom: Side view class average of N=80 single particles (Fig. S12, right). Scale bar: 25 nm.

We computed 2D class averages from single particle micrographs (Figure S12), which resulted in two distinct images (Figure 2b), corresponding to front and side view transmission projections of the particles. The front view displays four distinct layers of double-stranded DNA. The asymmetric feature is visible on the outer two helices on the top side (Figure 2a top). The class averaged image of the side view shows less detail of the expected 4 layers of DNA and the asymmetric feature. This is likely due to the staining with uranyl-formate and the tendency of the structures to adhere to the grid surface predominantly with their front side up resulting in less side-view particles. From these images, we measured an inner diameter of ~35 nm and a thickness of ~11 nm (Fig.S13). Accounting for the K_10_-PEG_5K_ coating by including a layer with thickness 2.4 ± 1.3 nm, as measured by Ponnuswamy et al.^21^ would result in an effective inner diameter of ~30 nm, and an effective height of ~16 nm including the unstructured PEG brushes. Class-averaged images of K_10_-PEG_5K_-coated and bare octagons match well (Figure S14), which shows that the coating preserves the global shape of the octagon pore.

### DNA origami pore reconstitution in liposomes by cDICE

We first tested spontaneous insertions observed for our 55 nm (outer diameter) large DNA origami pores into free-standing planar lipid bilayers or into pre-formed GUVs, under various conditions and with different nanopore versions with either 32 or 64 cholesterols. As expected, these attempts were unsuccessful. While membrane interactions drove the pores towards the GUVs (Fig.S16), there was a total lack of influx of molecules (Fig.S17), indicating that no pores were created with a functional transmembrane opening.

We therefore used an alternative approach where we incorporate the DNA origami pores during the process of the formation of the GUVs. We employed an inverted emulsion technique called cDICE^18^ which yields unilamellar liposomes with good encapsulation of a large variety of macromolecules^22^. Briefly, in cDICE, layers of buffer solution and lipid-in-oil suspension are deposited subsequently in a rotating chamber. As illustrated in Fig.3a, an inner buffer solution containing the origami pores is then injected through a capillary into the rotating lipid-in-oil suspension at a constant flow rate. Shear forces due to the rotation result in detachment of droplets from the capillary orifice. While travelling through the oil phase, these droplets acquire a first monolayer of lipids where the origami pores can insert. A second monolayer is subsequently formed when the droplets cross the oil-water interface, resulting in GUVs that are eventually collected from the outer buffer solution. In this cDICE process, we thus aimed to reconstitute the pore within the lipid monolayer that is formed in the water-in-oil droplets during the first step of cDICE. Whereas the origami rings with 64 cholesterols mainly resulted in formation of large aggregates at the droplet interface (Fig.S18), the rings with 32-cholesterols appeared to interact with the lipid monolayer without inducing significant aggregation, suggesting a more promising route. Hereafter, we report the results for the 32-cholesterols version of the DNA origami rings.

**Figure 3:**
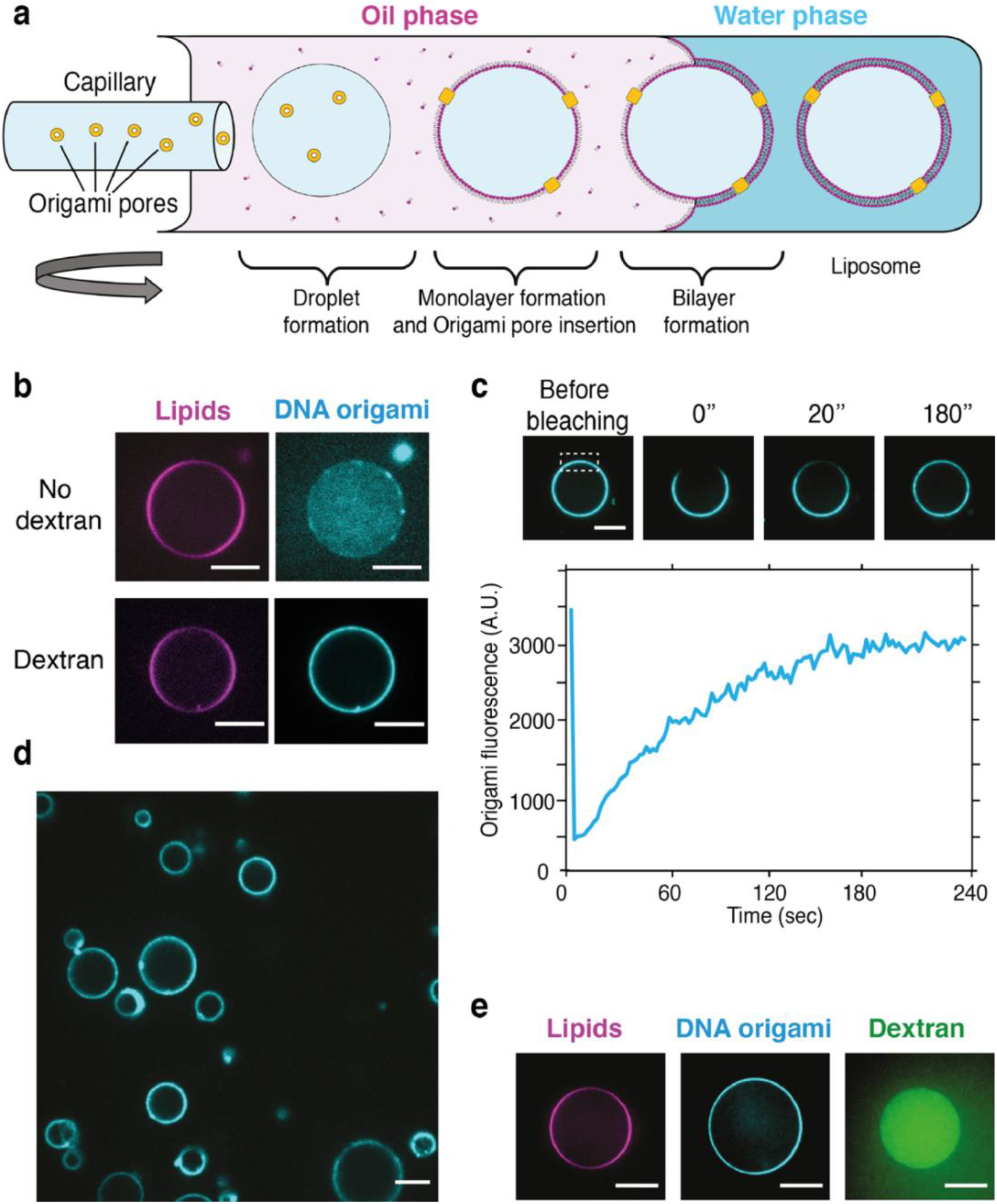
Reconstitution of wide DNA origami pores in GUVs by cDICE. **a,** Schematics of the cDICE workflow. A section of the rotating chamber is depicted, including the oil and outer buffer phases. Origami pores are indicated in yellow. **b,** Comparison of the efficiency of DNA origami pore localization at the membrane by addition of 2MDa dextran in the inner buffer. Without dextran, origami pores were observed to remain dispersed in bulk, whereas inclusion of dextran drove the origami pores to the lipid surface. **c,** FRAP analysis of origami pores reconstituted in GUV that shows that the pores are mobile within the lipid bilayer. The dashed rectangle indicates the bleached area, which recovers within a few minutes. **d,** Example of large field of GUVs with reconstituted origami pores. **e,** Example of a GUV retaining the 2MDa dextran in the lumen after 24 hours incubation in buffer. Scale bars: 10μm.

In a standard cDICE protocol^22^, the inner buffer solution that is injected contains either sucrose or Optiprep^22^ to obtain an optimal density of the inner GUV solution. We found, however, that in these conditions the reconstitution of 32-cholesterols origami rings was inefficient, with most of the origami pores remaining in solution after vesicle formation (Fig.3b). When we included high molecular weight dextran (~2 MDa) in the inner buffer, we found that 32-cholesterol origami rings could be fully localized at the membrane (Fig.3b). Note that the choice of such a large dextran molecule (2 MDa) was also motivated by the high diameter of gyration (~76 nm; Ref.^23^) which makes it large enough to stay in the vesicle despite the presence of the 30 nm wide DNA origami pores.

The origami pores appeared to be evenly distributed on the GUV surface, only occasionally forming clusters, which then were also enriched in lipids (Movie 1 and Movie 2: z-stack and 3D rendering). Moreover, Fluorescence Recovery After Photobleaching (FRAP) analysis revealed that DNA origami pores could freely diffuse within lipid bilayer of the vesicle membrane (Fig.3c). Hundreds of GUVs could be obtained from a single preparation (Fig.3d). The resulting GUVs retained the 2 MDa dextran over prolonged periods of time (>24hrs, Fig.3e) and were remarkably stable, remaining spherical in spite of osmotic changes, suggesting that the origami pores were very well able to equilibrate osmotic differences by allowing osmolytes to rapidly cross the membrane. All in all, the data indicate the successful insertion of wide DNA origami pores in the membrane of GUVs.

### Protein influx through the pores demonstrates their transmembrane transport capabilities

To assess whether the DNA origami pores were functional, i.e., inserted in the intended orientation that allows for transmembrane transport through the central cavity of the octagon, we tested the influx of IBB-mEGFP-Cys, a 34.6 kDa variant of the soluble green fluorescent protein that includes an importin β binding domain^24^ (see supporting information S22), henceforth referred to as ‘GFP’ for simplicity.. We measured the normalized fluorescence intensity difference I_n,diff_ = (I_out_−I_in_) / I_out_ of the vesicles, i.e., the difference between in-vesicle intensity (I_in_) and the outer bulk (I_out_) divided by I_out_ (Fig.4a) after overnight incubation of the GUVs in an outer solution with 2.3 μM of GFP. While pore-less vesicles (control, Fig.4b) remained empty yielding a finite value I_n,diff_ = 0.13 ± 0.05 (errors are S.D., N=34), the pore-containing liposomes showed influx of GFP for about 50% of the vesicles (Fig.4c): Half of the vesicles exhibited an almost complete saturation to I_n1,diff_ = 0.01±0.03 (N=45), while the other half yielded I_n2,diff_ = 0.09±0.03 (N=45), i.e., no significant change compared to the control. We note that heterogeneity in GUV samples is a widely reported phenomenon^25^, that here apparently is associated with variations in the efficiency of pore insertion during vesicle formation in cDICE. Most importantly, a clear fraction of the liposomes unambiguously showed transmembrane transport of the folded proteins, indicating transport through the ultrawide DNA origami pores.

**Figure 4.**
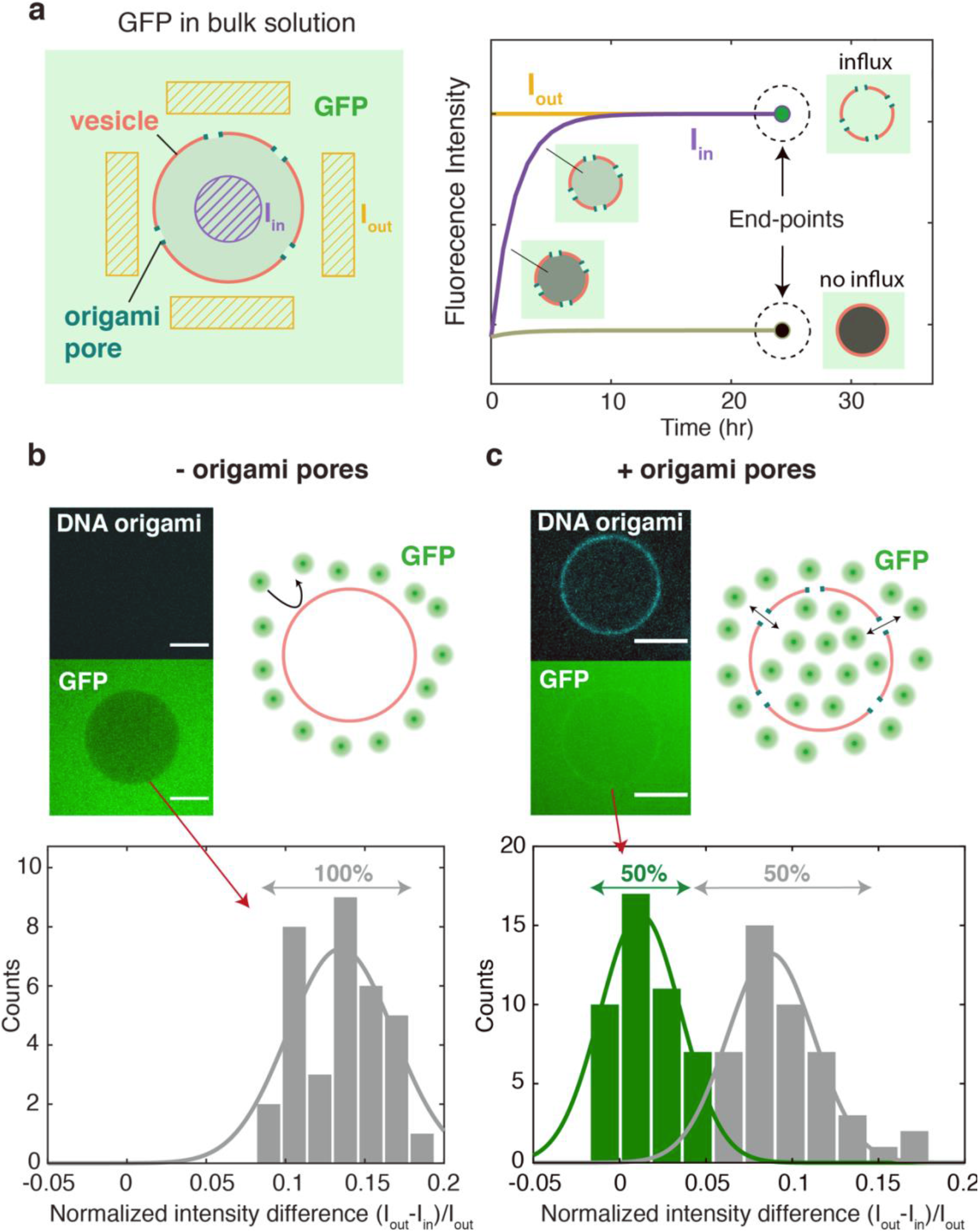
Test of GFP influx through origami pores. **a,** Schematic of the influx experiment. Left: Sketch of a vesicle (red) containing pores (blue), that is incubated with GFP (green). Right: Fluorescence intensities I_in_ and I_out_ that are expected to be measured in a time-lapse experiment on the dashed purple and yellow areas, respectively. The right part of the graph illustrates the end-point intensities after ~24hrs, after an initial transient. **b,** Influx experiment with vesicles that were not subjected to DNA-pore insertion. Top right: sketch of a pore-less vesicle that excludes GFP. Top left: example of fluorescence image for a pore-less vesicle (cyan) that prevents GFP (green) from entering the vesicle, showing up as a darker area signifying the absence of GFP. Bottom: histogram of the normalized end-point intensity difference (I_out_-I_in_)/I_out_ for the pore-less vesicles, verifying that no vesicles showed GFP influx. **c,** Influx experiment with vesicles that were functionalized with origami pores. Top right: sketch of a pore-containing vesicle filled with GFP. Top left: example of fluorescence image for a pore-containing vesicle (cyan) that allows GFP to diffuse (green) into the vescicle, resulting in I_out_-I_in_≈0. Bottom: histogram showing normalized end-point intensity difference (I_out_-I_in_)/I_out_ for the pore-less vesicles. We find that ~50% of the vesicle population manifested GFP influx. Scale bars: 10μm.

Next, we characterized the GFP influx rate quantitatively using a FRAP assay (Fig.5a). In this experiment, we selected pore-containing vesicles that showed that influx occurred during the overnight incubation (i.e., those with I_n,diff_ ≈ 0), then photobleached the inner part, and subsequently measured the recovery of the fluorescence signal. An example of FRAP images is shown in Fig.5b (top). Starting from an equilibrium state where I_in_≈I_out_ (Fig.5b, left), the vesicle fluorescence drops upon photobleaching to a level comparable to the one of an empty vesicle, to subsequently fully recover within a few minutes (Fig.5b, right). Notably this timescale is much faster compared to those of previously reported fluorescence recovery with DNA origami pores^7,8^. A control with vesicles lacking pores (Fig.5b, bottom) resulted in no appreciable recovery upon photobleaching. Figure 5c displays I_in_ and I_out_ for a pore-containing GUV as a function of time where, after initial transient behavior (grey area), I_out_ is constant, whereas I_in_ continues to slowly increase over time, which represents the influx of GFP through the ultrawide DNA origami pores.

**Figure 5.**
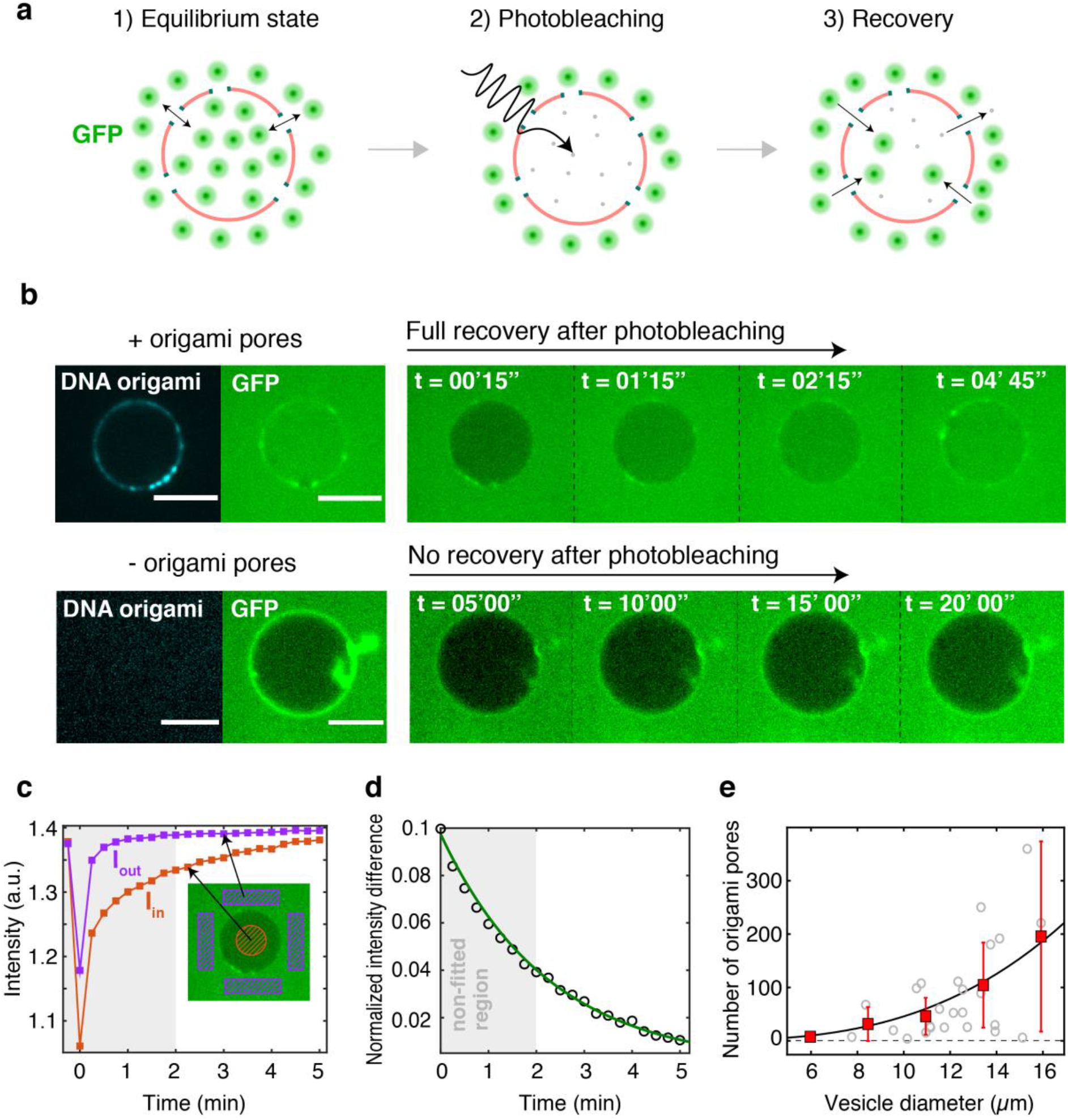
FRAP assay, modelling, and extraction of number of functional pores. **a,** Schematic of the FRAP assay: 1) the vesicle is left to equilibrate with bulk solution containing GFP; 2) GFP molecules inside the vesicle are photobleached; 3) new GFP molecules are driven through the ultrawide origami pores from the outer solution, causing a recovery of the fluorescence signal. **b,** Example of a FRAP experiment. Top: (left) image showing the pore-containing vesicle (origami pores in cyan) at equilibrium with the outer GFP (green) solution. Right: series of frames showing recovery (within minutes) of the GFP signal after photobleaching. Bottom: (left) image showing a pore-less vesicle (cyan) that excludes GFP (green) present in the outer solution. Right: series of frames showing the absence of a signal increase after photobleaching. **c,** Fluorescence intensities I_in_ (red) and I_out_ (purple) that reduce and recover after photobleaching. Grey area denotes the initial transient, after which I_out_ can be considered constant. Inset: highlight of the regions where I_in_ (red) and I_out_ (purple) were measured. **d,** Normalized intensity difference I_n,diff_ over time. Data are fitted by Equation (4). Grey area was excluded from the fit as the model assumes a constant I_out_. **e,** Number of origami pores extracted by fits for different vesicles, as a function of vesicle diameter (grey circles). Averaged data (red squares, error bars are S.D.) showed an increase of the number of pores as a function of vesicle diameter. Black line represents a cubic fit to the data. Scale bars: 10μm.

To obtain the number of functional pores, we model the influx rate using a diffusion model derived from the Fick’s first law^26^

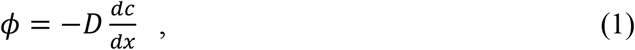

which describes the flux *ϕ* of GFP molecules with diffusion constant *D*, as a result of the concentration gradient 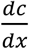 between the outer (*C*_*out*_) and inner (*C*_*in*_) environment of the GUV.

The flux *ϕ* is defined as number of molecules *dN* crossing a membrane area *dA* in a time *dt*. Since the vesicle volume *V* is constant over time, the in-vesicle concentration *C*_*in*_(*t*) is a time-dependent variable that reflects the fact that the vesicle is filling up over time, i.e., 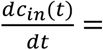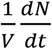. The outside concentration *C*_*out*_ instead is assumed to be constant over time, which is reasonable given the large volume (~200 μL) of the bulk solution as compared to the ~pL volume of the vesicle. Assuming that the diffusion occurs through a number *N*_*p*_ of DNA origami pores that each have a length *L*_*p*_ and an area *A*_*p*_, yielding a total integrated area *A* = *A*_*p*_*N*_*p*_, we can rewrite (1) as the first order differential equation

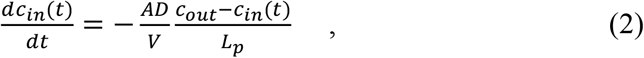

which, with *C*_*in*_(*t* = 0) = *C*_*in,start*_, has a solution

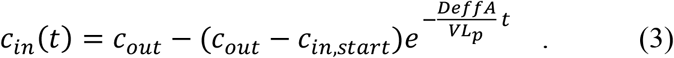

Note, however, that for the transport through the pores, we need to employ an effective diffusion constant *D*_*eff*_ which is reduced compared to the bulk diffusion constant *D* to account for the confined transport through the pores, as reported by Dechadilok and Deen^27^. For our pore dimensions, this results in almost a factor of 2 decrease of the GFP diffusivity, i.e., *D*_*eff*_ = 0.54 *D*_*bulk*_. Upon rearranging Equation (3), we then finally obtain

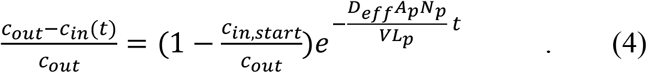

As the measured fluorescence intensities are a good measure for the protein concentrations of the fluorescent GFP, we can use Equation (4) to fit the I_n,diff_ (t) data, with 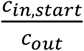 and *N*_*p*_ as fit parameters. Figure 5d provides an example, showing an excellent fit to the FRAP data. From fitting FRAP curves acquired from different vesicles, we find that the number of pores *N*_*p*_ varies between a few to a few hundred per GUV. As Fig.5e shows, *N*_*p*_ increases nonlinearly with the vesicle diameter. Assuming that all origami pores would move from the volume into the membrane during GUV formation, *N*_*p*_ should scale with the cube of the vesicle diameter, which is consistent with the behavior observed (black line in Fig.5e).

### The ultrawide DNA origami pores act as a molecular sieve with ~ 30 nm cut off

To provide evidence that our DNA origami pores indeed form open channels with an effective size of the order of the inner pore diameter of 30 nm, we tested the influx of dextran-FITC molecules with a variety of sizes, viz, D_g_ ≈ 15 nm (70kDa), 22 nm (150kDa), and 76 nm (2MDa), see Fig.6a, where Dg is the diameter of gyration as measured by Hanselmann and Burchard^23^. As in the GFP influx experiment, we incubated the pore-containing vesicles with 2μM of dextran-FITC, and measured the normalized intensity difference I_n,diff_ after ~24hrs. This was done independently for each of the 3 molecules, keeping the same molar concentration of dextran-FITC molecules in bulk solution. Similar to the GFP experiments (Fig.4c), I_n,diff_ of pore-containing vesicles yielded two populations when incubated with 70kDa dextran-FITC (Fig.6b, left), where 58% manifested influx (I_n1,diff,70k_= 0.02±0.03, errors are S.D., N=47), while 42% remained empty (I_n2,diff_,70k=0.13±0.02, N=34). Incubation with 150kDa dextran-FITC (Fig.6b, center) led to a similar result, with 47% GUVs that were filled (I_n1,diff,150k_=0.03±0.03, N=24), against 53% of empty vesicles (I_n2,diff, 150k_=0.11±0.03, N=27).

**Figure 6.**
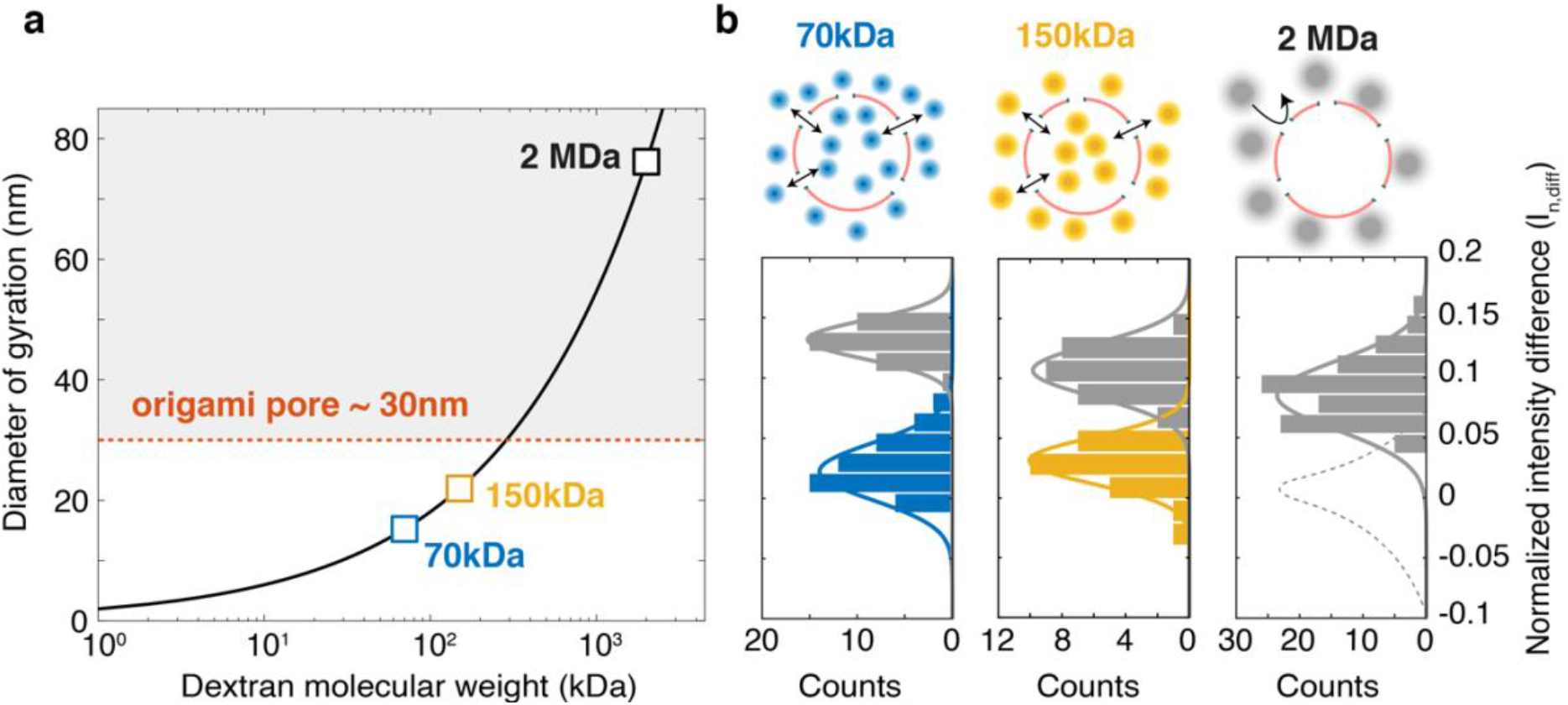
Size selectivity of DNA origami nanopores. **a,** Diameter of gyration of dextran *D*_*g*_ vs molecular weight *M*_*w*_. Black line is a fit of *D*_*g*_ = 0.072 ∙ *M*_*w*_0.48(Ref.^23^). Dextran sizes of 70 kDa, 150kDa, and 2MDa employed in our study are highlighted in blue, yellow, and black, respectively. Grey area underlies dextran sizes larger than the origami pore (red) which are not expected to translocate. **b,** Histograms showing normalized intensity difference of pore-containing vesicles after overnight incubation with 70kDa (left), 150kDa (center), and 2MDa (right). Blue and yellow bars indicate vesicles that showed influx of 70kDa and 150kDa, respectively, whereas grey bars represent empty vesicles. Dashed line in the rightmost histogram represents a hypothetical population of vesicles that would have been present if influx of 2MDa through the origami pores had occurred.

Importantly, this demonstrates that very large (150 kDa; 22 nm) macromolecules can be transported across the ultrawide DNA origami nanopores. By contrast, incubation with the even larger 2MDa dextran-FITC (Fig.6b, right) resulted in only one population of empty vesicles, with I_n,diff,2M_=0.09±0.04 (errors are S.D., N=98), indicating a lack of any transmembrane transport during 24 hours. This is fully consistent with expectations as its size clearly exceeds the 30 nm size of the origami pore. It is also consistent with its use as a macromolecular crowder, cf. Fig. 3e. The slight increase in background fluorescence (causing I_n2,diff,70k_> I_n2,diff,150k_> I_n2,diff,2M_) can be attributed to the fact that larger dextran-FITC molecules possess a higher number of FITC fluorophores (see ‘Methods’ section).

## Discussion and conclusion

In this work, we achieved functional reconstitution of ultrawide DNA origami pores in giant liposomes, obtaining stable transmembrane pores of 30 nm inner diameter within the lipid bilayer. We successfully achieved the pore reconstitution by inserting of the origami pore into a lipid monolayer followed by bilayer formation. This methodology lowers the energy barrier for pore insertion, allowing functional reconstitution of pores with much larger diameter than previously possible.

Our results open the way to a number of biomimetic applications. For example, future efforts can be directed towards the functionalization of the origami pore with FG-nucleoporins from the Nuclear Pore Complex (NPC), the macromolecular complex mediating transport across the nuclear envelope in eukaryotes^28^, as reported in previous works^14,15^. This will enable the creation of selective biomimetic pores that permit the exclusive transport of specific transport receptor proteins. While this constitutes an exciting platform for investigating nucleocytoplasmic transport *in vitro*, it may also be employed as a building block for synthetic cells, thus paving the way towards the *in vitro* reconstitution of an artificial model of a nucleus. This will, for example, aid the study of emergent properties of liposome-confined genomes^29^, by allowing the controlled injection of genome-processing proteins. Indeed, our DNA origami pores may be a great asset for building a synthetic cell as they provide a stable portal that is large enough for biological macromolecules like proteins and RNAs to enter a liposome. The long-term stability and functionality of these liposomes is a particularly useful feature in such synthetic-biology applications. Finally, responsive drug-delivery systems that release clinically relevant macromolecules such as antibodies upon stimulation by specific biomolecules could be envisioned as well. We expect that the ultrawide DNA origami nanopores may thus find a broad variety of different applications.

## Acknowledgements

We thank Kristina Ganzinger and Gijsje Koenderink for helpful discussions and technical advice in setting up cDICE; Federico Fanalista, Nils Klughammer, Jaco van der Torre for discussions; Jeremie Capoulade for technical support on confocal microscopy; and Dimitri de Roos for constructing the cDICE steup. This work was supported by a European Research Council Consolidator Grant to H.D. (GA no. 724261), the Deutsche Forschungsgemeinschaft through grants provided within the Gottfried-Wilhelm-Leibniz Program and the SFB863 Project ID 111166240 TPA9 (to H.D.) C.D. acknowledges funding support from the ERC Advanced Grant 883684, NWO grants 16PR3242-1, BBOL.737.016.016, and OCENW.GROOT.2019.068, and the NanoFront and BaSyC programs.

## Author contributions

A.F., H.D, and C.D. conceived the research. N.D.F. and C.D. set up cDICE and devised the reconstitution protocol. P.S. and H.D. devised the DNA origami nanopores. P.S. carried out assembly, quality control, and TEM imaging of the DNA origami nanopores. A.F. and N.D.F carried out influx experiments. A.F. performed analysis and modelling of influx data. E.O.v.d.S. expressed and purified the GFP. A.F., N.D.F., P.S., H.D., and C.D. wrote the manuscript.

## Methods

### Design of scaffolded DNA origami objects

The objects were designed using caDNAno v0.2^30^.

### Optimal folding conditions of the octagon

The folding reaction mixture for the octagon contained scaffold DNA^31^ at a final concentration of 10 nM and oligonucleotide strands (IDT Integrated DNA Technologies) at 300 nM each. The folding reaction buffer contained 5 mM TRIS, 1 mM EDTA, 5 mM NaCl (pH 8) and 24 mM MgCl_2_. The folding reaction mixtures were subjected to the following thermal annealing ramp using TETRAD (MJ Research, now Biorad) thermal cycling devices: 15 minutes at 65°C, followed by 16 hours at 50°, followed by incubation at 20°C before further sample preparation steps.

### Purification of DNA origami nanostructures

Excess oligonucleotides were removed via ultrafiltration (Amicon Ultra 0.5 ml Ultracel filters, 50K) with FoB5 (5 mM TRIS, 1 mM EDTA, 5 mM NaCl and 5 mM MgCl_2_) buffer19. All centrifugation steps were performed at 10 kG for 3 minutes at 25°C. Step 1: 0.5 ml of FoB5 buffer was added to the filters and centrifuged. Step 2: 0.1 ml of folded object sample and 0.4 ml of FoB5 buffer was added to the filters and centrifuged. Step 3: 0.45 ml FoB5 buffer was added to the filters and centrifuged. Step 3 was repeated 3 times before a final retrieving step: the filter insets were turned upside down, placed into a new tube and centrifuged. This concludes one round of ultrafiltration.

### Coating of the octagon

Different volumes of purified octagons were mixed with PEG-oligolysine (K_10_-PEG_5K_). The optimal amount of K_10_-PEG_5K_ added was determined based on the ratio of nitrogen in amines to the phosphates in DNA (Ponnuswamy et al.^21^), to a final ratio of 0.75:1 (N:P).

### Negative-stain TEM

A carbon Formvar grid (Electron Microscopy Sciences) was plasma-treated (45 seconds, 35 mA) right before use. 5 μl of uncoated octagon sample (10 - 50 nM DNA object concentrations, 5 – 20 mM MgCl_2_ concentration) was pipetted onto the grid. Coated octagon samples were pipetted onto the grids without plasma-treatment, in order to reduce positive staining effects. The sample droplets were incubated for 30 - 120 s on the grids and then blotted away with filter paper. A 5 μl droplet of 2% aqueous uranyl formate (UFO) solution containing 25 mM sodium hydroxide was pipetted onto the grid and immediately blotted away. A 20 μl UFO droplet was then pipetted onto the grid, incubated for 30 s and blotted away. The grids were then air dried for 20 minutes before imaging in a Philips CM100 and an FEI Tecnai 120 microscope. Particles were picked automatically using crYOLO^32^ and subsequently class averaged using Relion3.0^33^.

### Gel electrophoresis of octagon samples

Octagon samples were electrophoresed on 1.5 and 2.0% agarose gels containing 0.5× tris-borate-EDTA and 5 mM MgCl_2_ (unless noted otherwise) for around 1-2 hours at 70 or 90 V bias voltage in a gel box, cooled in a water or ice water bath. The loaded samples contained final monomer concentrations of 10 nM (unless otherwise noted). The gels were scanned with a Typhoon FLA 9500 laser scanner (GE Healthcare) at a resolution of 25 or 50 μm/pixel. Further enhancing resulting images or parts of images was performed using Photoshop CS5.

### DNA origami sample preparation protocol

1. Folding of DNA origami samples at previously described optimal folding conditions.
2. Three rounds of ultrafiltration to purify the samples of excess oligonucleotides.
3. Addition of Atto647N modified oligonucleotide at a final oligonucleotide:DNA origami ratio of 48:1 (12 handles per DNA origami; effective 4:1 ratio).
4. One round of ultrafiltration to purify the samples of excess Atto647N oligonucleotides.
5. Addition of PEG-oligolysine at N:P ratio of 0.75:1, incubate at room temperature overnight.
6. Addition of cholesterol modified oligonucleotide at a final oligonucleotide: DNA origami ratio of 128:1 (32 handles per DNA origami; effective 4:1 ratio).
7. One round of ultrafiltration to purify the samples of excess cholesterol oligonucleotides.

### Reagents

The 2MDa dextran (D5376), MgCl_2_ (M8266), glucose (G7021), Optiprep (60% (w/v) iodixanol in water, D1556), Silicon oil (317667) and Mineral oil (M3516-1L) were purchased from Sigma Aldrich. Tris-HCl (10812846001) was purchased from Roche. DOPC (1,2-dioleoyl-sn-glycero-3-phosphocholine) (850375) and DOPE-Rhodamine (1,2-dioleoyl-sn-glycero-3-phosphoethanolamine-N-(lissamine rhodamine B sulfonyl) (ammonium salt)) (810150C) were purchased from Avanti Lipids. Lipids were stored and resuspended in anhydrous chloroform (288306, Sigma Alrich). Sodium dodecyl sulfate (SDS) employed for cleaning steps was purchased from Sigma Aldrich. UltraPure Bovine Serum Albumine (BSA) used for passivation of the glass coverslips was purchased by ThermoFisher. For the influx experiments, we employed 70kDa dextran-FITC (FITC:Glucose (mol/mol) = 0.004) (46945, Sigma Aldrich), 150kDa dextran-FITC (FITC:Glucose (mol/mol) = 0.004) (74817, Sigma Aldrich), and 2MDa dextran-FITC (FITC:glucose (mol/mol) = 0.001-0.004) (FD2000S, Sigma Aldrich).

### Expression and purification of IBB-eGFP

For the expression of His14-TEV-IBB-mEGFP-Cys (‘IBB-GFP’) plasmid pSF807 (Eisele et al.^34^) was transformed into *Escherichia coli* BLR cells. Cultures were grown in TB medium supplemented with 50 μg/ml kanamycin, and expression was induced with 0.5 mM IPTG during overnight incubation at 18°C, shaking at 150 rpm in baffled flasks. Cells were harvested by centrifugation (10 min 4000 rpm, JLA8.1000 rotor), washed in 1x PBS buffer and lysed using a French Press (Constant Systems) at 20 kpsi at 4°C in buffer A (2 M NaCl, 50 mM Tris/HCl pH 8.0, 5 mM MgCl_2_). Unbroken cells, debris and aggregates were pelleted in a Ti45 rotor (30 min, 40.000 rpm, 4°C), and the lysate resulting from 1 liter cell culture was applied to 2 mL (bed volume) of Ni-NTA matrix pre-washed with buffer A. After 60 min incubation while slowly inverting the tube in a cold room (4°C) the matrix was washed extensively with buffer A supplemented with 60 mM imidazole, and buffer B (50 mM Tris/HCl pH 7.5, 300 mM NaCl, 5 mM MgCl_2_, 1 mM βME, 10% glycerol). Following overnight elution by homemade TEV protease in buffer B, the eluate was concentrated by ultracentrifugation and further purified by size exclusion chromatography on a Superdex 200 column (GE Healthcare) pre-equilibrated with buffer B supplemented with 0.5 mM EDTA. The final IBB-GFP sample with a concentration of 138 μM was aliquoted, snap frozen in liquid nitrogen and stored at - 80°C.

### cDICE buffers and solutions

The encapsulated inner solution contains 50mM Tris-HCl pH 7.4, 5mM MgCl_2_, 15μM 2MDa dextran, 2 nM DNA origami pores. The outer solution contains glucose in milliQ water and is osmotically matched to the inner solution. The oil phase is a mixture of silicone and mineral oil (4:1 ratio) and contains lipids dissolved at 0.2mg/mL. The lipid-in-oil suspension was prepared as in Van de Cauter et al.^22^ and used immediately.

### Preparation of lipid-in-oil suspension

DOPC and DOPE-Rhodamine lipids solubilized in chloroform were mixed at 99.9:0.1 ratio and blow-dried with pure nitrogen. Inside a glove-box filled with pure nitrogen, lipids were subsequently resolubilized with anhydrous chloroform. The freshly prepared mixture of silicon and mineral oil was added to the lipids dropwise while vortexing at 1400 rpm. The lipid-in-oil solution was finally vortexed at 2800 rpm for two minutes, further sonicated in an ice bath for 15 minutes.

### cDICE equipment and GUV production

The rotating component of the cDICE device is built from a Magnetic stirrer L-71 (LAbinco). A Teflon adaptor connects the rotating shaft with the cDICE chamber, assembled from a standard 3.5 cm cell culture dish (corning) modified to decrease the height. The chamber has an inner height of 8 mm and a circular opening of 10 mm on the top to allow insertion of the capillary and collection of the GUVs. The detailed protocol for GUV production is reported in Van de Cauter et al.^22^, while here we describe it briefly. First, the cDICE chamber is set to rotate at 300 rpm. 350 μL of outer aqueous solution is injected into the rotating the chamber, followed by 1 mL of lipid-in-oil solution, resulting in the two phases being stacked as depicted in Fig.3a. The tip of a 100 μm PEEK capillary tube (211633-3, BGB) is then inserted into the oil phase of the rotating chamber, allowing for a continuous supply of inner aqueous solution into the oil phase by a pressure pump (MFCS-EZ, Fluigent) set at 900 mbar. After 15 min, the rotating speed is set to zero, and GUVs are collected from the outer aqueous solution and moved to a pre-passivated imaging well to carry out the fluorescence experiments.

### Data collection and analysis

Fluorescence images were acquired using spinning disk confocal laser microscopy (Olympus IXB1/BX61 microscope, 60x objective, iXon camera) with Andor iQ3 software. To induce GFP photobleaching, we employed raster scanning 491 nm laser (at 9.8 mW) over the region of interest. To measure the recovery signal, frames were collected every 15 seconds, starting right after the photobleaching event. Fluorescence images were analyzed and processed using ImageJ (v2.1.0). The extracted fluorescence data were plotted and fitted using MATLAB.

### Diffusion model

Fluorescence recovery data were fitted with a diffusion model, described by Equation (4), derived from the Fick’s law of diffusion. While 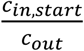 and *N*_*p*_ were free parameters of the fit, all the other parameters were known or could be calculated. More particularly, the area of the pore *A*_*p*_ = 745.6 nm^2^ was calculated from assuming an octagonal pore with an effective radius of 15 nm (as discussed in the text). The effective diffusion constant *D*_*eff*_ = 47 μm^2^/s of GFP was calculated using the formula provided by Dechadilok and Deen^27^

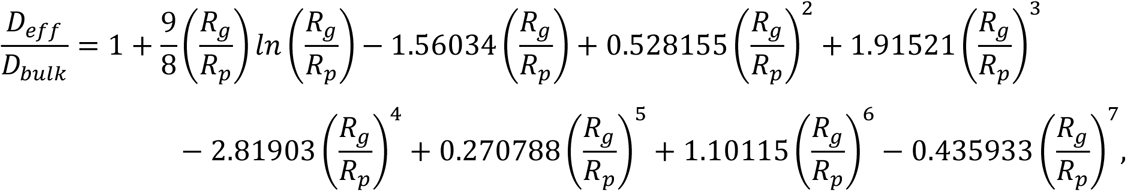

where *R*_*g*_ is the molecule radius of gyration (for GFP, *R*_*g*_ = 3.5 nm), *R*_*p*_ is the pore radius (here 15 nm), and *D*_*bulk*_ is the diffusion constant of the molecule in free solution (for GFP, *D*_*bulk*_ = 87 μm^2^/s, Ref.^35^). The channel length *L* = 16 nm was assumed to equal the effective height of the origami, which was measured by TEM imaging (Fig.S13b). Finally, *V* was the volume of the vesicle which was estimated from the fluorescence images.

